# A matter of attention: Crossmodal congruence enhances and impairs performance in a novel trimodal matching paradigm

**DOI:** 10.1101/015958

**Authors:** Jonas Misselhorn, Jonathan Daume, Andreas K. Engel, Uwe Friese

## Abstract

A novel crossmodal matching paradigm including vision, audition, and somatosensation was developed in order to investigate the interaction between attention and crossmodal congruence in multisensory integration. To that end, all three modalities were stimulated concurrently while a bimodal focus was defined blockwise. Congruence between stimulus intensity changes in the attended modalities had to be evaluated. We found that crossmodal congruence improved performance if both, the attended modalities and the task-irrelevant distractor were congruent. If the attended modalities were incongruent, the distractor impaired performance due to its congruence relation to one of the attended modalities. Between attentional conditions, magnitudes of crossmodal enhancement or impairment differed. Largest crossmodal effects were seen in visual–tactile matching, intermediate effects for audio–visual and smallest effects for audio–tactile matching. We conclude that differences in crossmodal matching likely reflect characteristics of multisensory neural network architecture. We discuss our results with respect to the timing of perceptual processing and state hypotheses for future physiological studies. Finally, etiological questions are addressed.

## 1 Introduction

Any instant of conscious perception is shaped by the differential contributions of all of our sensory organs. Integration of these different sensory inputs is not merely additive but involves crossmodal interactions. The mechanisms of these interactions are still far from being completely understood. One on-going challenge in the field of multisensory research is the question of how crossmodal interactions can be identified and quantified (Gondan & Röder, 2006; Stevenson et al., 2014).

In many cases, crossmodal interactions have been investigated by means of redundant signal detection paradigms in which performance in unimodal trials is compared to performance in redundant multimodal trials (Diederich & Colonius, 2004). The race model inequality introduced by Miller (1982) is commonly used to decide if performance increments are actually indicative of crossmodal interactions. These are inferred if the multimodal cumulative distribution function (CDF) of response times is larger than the sum of the unimodal CDFs. Otherwise, performance increments are deemed to be due to statistical facilitation (Raab, 1962). Stimulus design and presentation in redundant signal detection paradigms were typically chosen such that multisensory principles formulated by Meredith and Stein (1993) could be tested. These principles were deduced from the observation that multisensory neurons showed strongest crossmodal effects when stimuli to distinct modalities shared temporal and spatial characteristics, and crossmodal effects increased when unimodal stimulus intensities decreased (Sarko et al., 2012).

Although supporting evidence for the validity of these principles in human behavior exists (e.g., Bolognini, Frassinetti, Serino, & Làdavas, 2004; Senkowski, Saint-Amour, Höfle, & Foxe, 2011), an increasing number of empirical null results and methodological issues question the general applicability of these principles (Holmes, 2007; Otto, Dassy, & Mamassian, 2013; Pannunzi et al., 2015; Spence, 2013; Sperdin, Cappe, & Murray, 2010). Additionally, it has been demonstrated that crossmodal interactions can also have competitive effects, leading to crossmodal inhibition rather than enhancement (Sinnett, Soto-Faraco, & Spence, 2008). In an audio–visual redundant signal detection paradigm, auditory detection was delayed by redundant visual presentation while visual detection was speeded by redundant auditory presentation. Similarly, Wang and colleagues (2012) presented results suggesting the coexistence of crossmodal inhibition and enhancement in a trimodal study. In a target detection task, participants were presented with visual, auditory or somatosensory targets in the presence or absence of perceptual competition by the respective other modalities. Overall, visual detection was fastest whereas auditory and somatosensory detection was comparable. Interestingly, they observed that the detection of auditory targets was impaired by perceptual competition while vision was unaffected and tactile detection facilitated. These crossmodal effects can be understood in the context of perceptual gain adjustments due to multisensory interactions (Yi Jiang & Shihui Han, 2007).

The inconsistency of results concerning crossmodal interactions might relate to aspects of multisensory processing that are typically not addressed by redundant signal detection paradigms. An important aspect of crossmodal interactions is the evaluation of crossmodal matching or conflict. As a result, matching sensory inputs will be bound into a coherent percept of an event, whereas sensory information will be processed separately if it is unlikely that a common origin is emitting these signals (Engel & Singer, 2001; Senkowski, Schneider, Foxe, & Engel, 2008; Treisman, 1996). Accordingly, congruence between stimulus features in distinct sensory modalities leads to crossmodal enhancement of sensory processing, resulting in effective binding of corresponding neural representations. By contrasting congruent to incongruent multisensory stimulus presentation, the extent of crossmodal congruence enhancement can be probed. Congruence in this context can be related to low-level spatiotemporal characteristics but also to more complex stimulus features such as, for example, semantic aspects of the stimuli or crossmodal correspondences. Concerning the former, detection of natural objects was improved if auditory and visual information matched semantically (Schneider, Debener, Oostenveld, & Engel, 2008; Yuval-Greenberg & Deouell, 2007). Crossmodal correspondences, on the other hand, relate to stimulus features that are consistently associated crossmodally and are, thus, described as “natural cross-modal mappings” (Evans & Treisman, 2010). A very robust correspondence, for instance, could be established between auditory pitch and visual brightness (Marks, 1987). That is, detection of visual and auditory stimuli is improved when jointly presented in a congruent fashion (e.g. high pitch tone and bright visual stimulus) compared to incongruent presentation (e.g. low pitch tone and bright visual stimulus). Relations between the mechanisms of crossmodal correspondences and synesthetic experiences are being discussed (Spence, 2011).

In addition to stimulus-driven aspects of multisensory integration, there is growing awareness that attention plays a central role in how multisensory inputs are being integrated (Talsma, 2015). One possibility is that attention-related top-down mechanisms may generally enhance sensory processing of an attended event at the expense of other sensory input (Desimone & Duncan, 1995; Kastner & Ungerleider, 2001; Wascher & Beste, 2010). The interplay between attention and mechanisms of crossmodal integration, however, is more complex and impact seems to be exerted mutually (Talsma, Senkowski, Soto-Faraco, & Woldorff, 2010). On the one hand, it was shown that spatial attention directly interfered with mechanisms of crossmodal integration in an audio–visual redundant target detection paradigm (Talsma & Woldorff, 2005). In this study, multisensory effects on event-related potentials at fronto-central sites were larger for attended stimuli compared to unattended stimuli. Available data suggest that attention might not modulate crossmodal interactions in an all-or-nothing manner, but rather shapes the nature or magnitude of crossmodal integration (De Meo, Murray, Clarke, & Matusz, 2015). Supporting evidence for this has also been obtained in studies where congruence-related crossmodal effects are larger when attention is divided between modalities compared to when it is focused on one modality (Göschl, Engel, & Friese, 2014; Mozolic, Hugenschmidt, Peiffer, & Laurienti, 2008). On the other hand, it was shown that attentional allocation can be driven by mechanisms of multisensory integration in a visual search paradigm (Van der Burg, Olivers, Bronkhorst, & Theeuwes, 2008). Reaching similar conclusions, van Ee and colleagues (2009) showed that crossmodal congruence enhanced attentional control in perceptual selection.

Here we propose a novel trimodal matching paradigm to investigate interactions between attention and crossmodal congruence. To that end, participants receive simultaneous visual, auditory, and somatosensory stimulation on each trial. All three stimuli undergo a brief, simultaneous change in intensity (either increase or decrease) resulting in varying patterns of crossmodal congruence across trials. Per block, an attentional focus including two relevant modalities is defined for which congruence has to be evaluated irrespective of the third, task-irrelevant modality. Four different congruence patterns can be discerned (see Fig. 1 for an example)

**Fig. 1.**
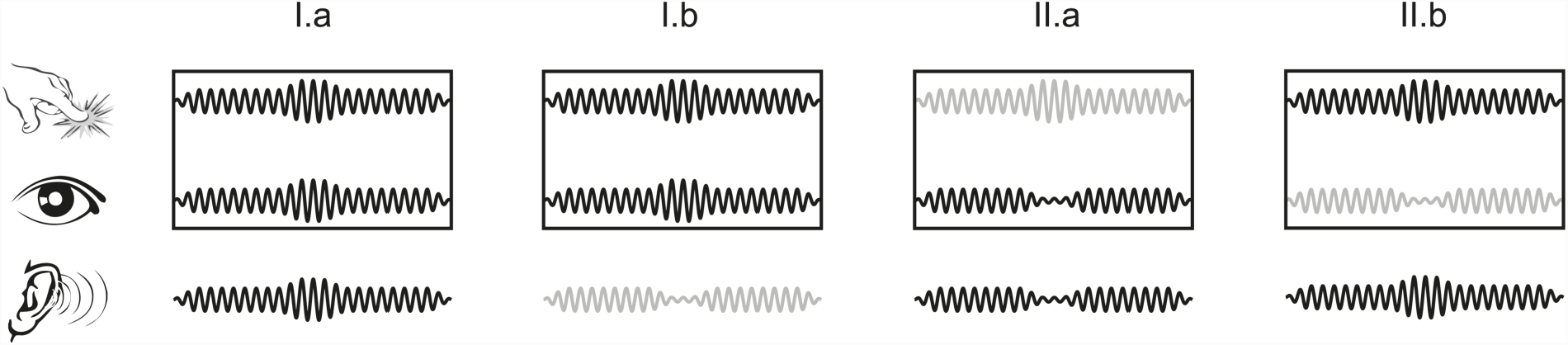
Congruence patterns for the visual–tactile focus (analogous for audio–visual and audio–tactile foci). Cartoons depict tactile (first row), visual (second row) and auditory (last row) stimulation. Boxes indicate the attended combination of modalities. Congruently changing modalities are depicted by a black line and incongruent modalities by a gray line. While all stimuli are congruent in I.a, one stimulus is deviant in I.b and II.a+b, respectively. In I.b, attended stimuli are congruent and the deviant is task-irrelevant. For the two attended incongruent cases, the auditory distractor is either congruent to the attended visual stimulus (II.a, audio–visual congruence) or to the attended tactile stimulus (II.b, audio–tactile congruence).

(I.a) All stimuli are congruent.

(I.b) The attended modalities are congruent, and the task-irrelevant modality is incongruent to both attended modalities.

(II.a) The attended modalities are incongruent, and the task-irrelevant modality is congruent to attended modality 1.

(II.b) The attended modalities are incongruent, and the task-irrelevant modality is congruent to attended modality 2.

In contrast to redundant signal detection paradigms, this trimodal paradigm requires the participant to evaluate crossmodal congruence and thereby ensures that information from both attended modalities must be processed in order to reach a decision. In a study using a similar paradigm, crossmodal congruence significantly speeded responses for audio-visual matching (Friese et al, in preparation). Employing a trimodal design allows to compare crossmodal matching between different bimodal foci of attention and to study the influence of the respective distractor. Our hypotheses are three-fold. First, in line with the literature on redundant target effects, we expect that performance in fully congruent trials is superior to trials with a task-irrelevant deviant (Fig. 1; I.a vs. I.b). Second, we hypothesize that the magnitude of performance will depend on how easily the distracting, task-irrelevant modality can be inhibited. This will in turn depend on the attentional focus. Based on Wang et al. (2012), it is assumed that vision is hardest to ignore and audition easiest. Accordingly, performance in the visual–tactile focus condition should be best while performance in the audio–tactile focus condition should be worst. Third, with respect to incongruent task-relevant stimuli, we expect that congruence between one attended stimulus and the distractor impairs performance. Mirroring the expectations outlined above, the distracting effect of congruence should also depend on the focus of attention.

## 2 Methods

### 2.1 Participants

Forty-nine participants were recruited for the study and received monetary compensation for their participation. Due to the high demands of the task, fifteen candidates were not able to complete the training successfully and were excluded from further participation (performance was below an average of 70% correct answers after 30 min of training). The remaining 34 participants were on average 24 years old (±4 years) and 20 of them were male (14 female). All had normal or corrected to normal vision and had no history of neurological or psychiatric disorders. After an explanation of the experimental procedure, participants gave written consent. The ethics committee of the Medical Center Hamburg – Eppendorf approved the study.

### 2.2 Stimulation

Participants were seated comfortably in a sound attenuated and dimly lit chamber. In all trials, three sensory modalities were stimulated by signals of 2 s that underwent a transient change in amplitude. Visual stimulation consisted of a drifting, continuously expanding circular grating presented centrally on a CRT screen (distance: 70 cm, visual angle: 5°) against a gray background. Phase onsets were randomized and contrast of the grating was experimentally increased or decreased. Auditory stimulation was delivered via ear-enclosing headphones (Ultrasone HIFI 680, transmission range 15 – 25.000 Hz). A complex sinusoidal sound was created by modulating high-frequency carrier signals (13 sine waves: 64 Hz and its first 6 harmonics as well as 91 Hz and its first 5 harmonics) with a low-frequency modulator (0.8 Hz) and was presented to the participants binaurally at 70 dB SPL. Changes of the complex tones were amplitude modulations with cosine-tapered transitions. Vibro-tactile stimulation was administered to both index fingers with C2 tactors (diameter: 2.97 cm, height: 0.76 cm, optimal stimulation frequency: 200 – 300 Hz, see http://www.atactech.com/PR_tactors.html). Two identical C2 tactors were embedded into custom-made foam retainers to prevent repulsion related modulation of the vibration. They were located on a tray attached to the chair ensuring a comfortable posture throughout the whole experiment. Vibro-tactile stimulation was set to a frequency of 250 Hz and the amplitude of the perpendicular contactor displacement was experimentally increased or decreased. Phase onset of the complex sound and vibro-tactile stimulus were locked to the phase onset of the expanding grating.

### 2.3 Experimental Paradigm

A central fixation dot was presented for 1000 – 1600 ms (see Fig. 2.a). Concurrent visual, auditory, and somatosensory stimulation lasted for 700 – 1000 ms until a change in stimulus intensity occurred for 300 ms simultaneously in all modalities (ramping up: 100 ms, maximal change intensity: 100 ms, ramping down: 100 ms). After change offset, stimulation continued for another 700 – 1000 ms depending on the pre-change interval length (total duration of stimulation was always 2 s). After presentation, participants responded verbally (response interval max. 3000 ms) and received feedback for 600 ms (ISI = [3600, 4200] ms). The participants’ task was to evaluate congruence between changes of an attended pair of modalities irrespective of the change in the third modality. The attentional focus was cued block-wise as audio–visual (AV), visual–tactile (VT) or audio–tactile (AT). A change was defined as “congruent” if the changes in the two attended modalities had the same direction, e.g., an increase in contrast of the grating and a concurrent increase in amplitude of the vibration while VT was cued for this block (Fig. 1, I.a+b). The direction of change in the ignored modality was irrelevant for the decision. A change was defined as “incongruent” if the directions of change in the attended modalities differed, e.g., an increase in vibration intensity and a concurrent decrease in contrast of the grating while VT had been cued for this block (Fig. 1, II.a+b).

**Fig. 2.**
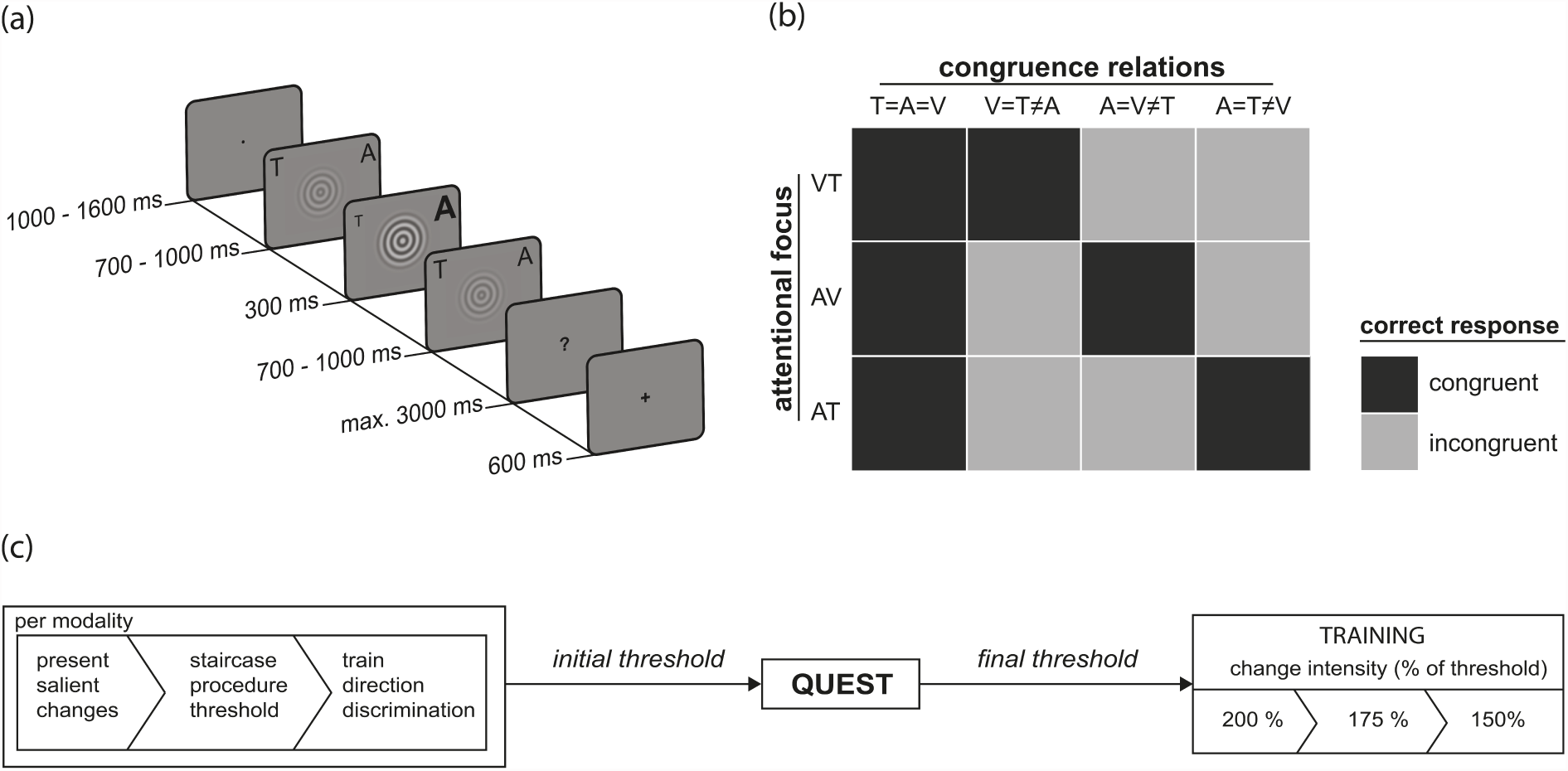
Overview of study design and procedure. In this example of a trial (a), visual contrast increases (central grating), vibration strength (T) decreases and auditory amplitude (A) increases. Correct answers (b) for all possible congruence relations (‘=’ is congruent, ‘≠’ is incongruent) per attentional focus (VT = visual-tactile, AV = audio-visual, AT = audio-tactile). Thresholding and training procedure (c) preceding the experiment.

After each trial, the participant was asked to report verbally if the change in the attended modalities was congruent (“gleich”, German for “equal”) or incongruent (“verschieden”, German for “different”). To exclude possible response time (RT) differences on the basis of differences in the vocalization of the two response options, participants initially said “Yes” as soon as they felt ready to respond. RTs were measured from change onset until verbal response onset (“Yes”). Subsequently, the response (“gleich”/”verschieden”) was given and evaluated online using custom-made speech recognition software. Feedback was given after each trial and at the end of each block. Each block consisted of 64 trials in random order and each stimulus configuration occurred 8 times. The order of attentional foci was randomized for each participant (e.g. AT, AV, VT) and repeated 5 times. The complete experiment comprised a total of 960 trials (15 blocks with each 64 trials). For each subcondition (cells in Fig. 2b), 80 trials were recorded.

### 2.4 Thresholding Procedure

The magnitude of change for a given modality and direction (increase or decrease) was individually adapted to ensure comparable perceptual salience and detection performance across modalities and change directions. The following procedure was implemented after extensive piloting (Fig. 2c). Initially, salient unimodal sample trials of both changes per modality were presented to get the participants acquainted with the stimulus material. These trials were repeated until the participants reported to have an intuitive feeling for how these stimuli and, most importantly, their changes look, sound or feel. Subsequently, a threshold was estimated with a reversed staircase procedure. Initial change magnitude was close to zero and was incremented from trial to trial with a fixed step-size. This was repeated until participants reported to be absolutely sure to have perceived the change. Subsequently, the direction of changes with the previously estimated magnitudes had to be judged by the participants. If they were not able to correctly categorize 80 % or more of the presented changes, estimated thresholds were increased and the task was repeated. After that, the actual threshold estimation was carried out using an implementation of the Bayesian adaptive psychometric method QUEST (Watson & Pelli, 1983). To this end, all three stimuli were presented concurrently with only one stimulus changing per block that had to be attended. The occurrence of increases and decreases of intensity was randomized and the magnitude of change was iteratively varied over 30 trials per change direction (see Fig. 3). The estimate from the preceding staircase procedure served as the initial prior. Subsequently, the last trial from the respective change direction in combination with information on success or failure to detect the change served as input for the Bayesian model to compute presentation intensity for the next trial. After 30 trials, stable estimates of detection thresholds were reached. The subsequent training comprised three levels. At the first level, change magnitudes were set to 200 % of the estimated threshold to facilitate task learning. When all conditions were mastered at a performance level of 70 % correct answers or more, the second level with changes at a magnitude of 175 % of the estimated threshold followed. Finally, changes were presented at 150 % of the estimated threshold. This magnitude of changes was used in the experimental blocks.

**Fig. 3.**
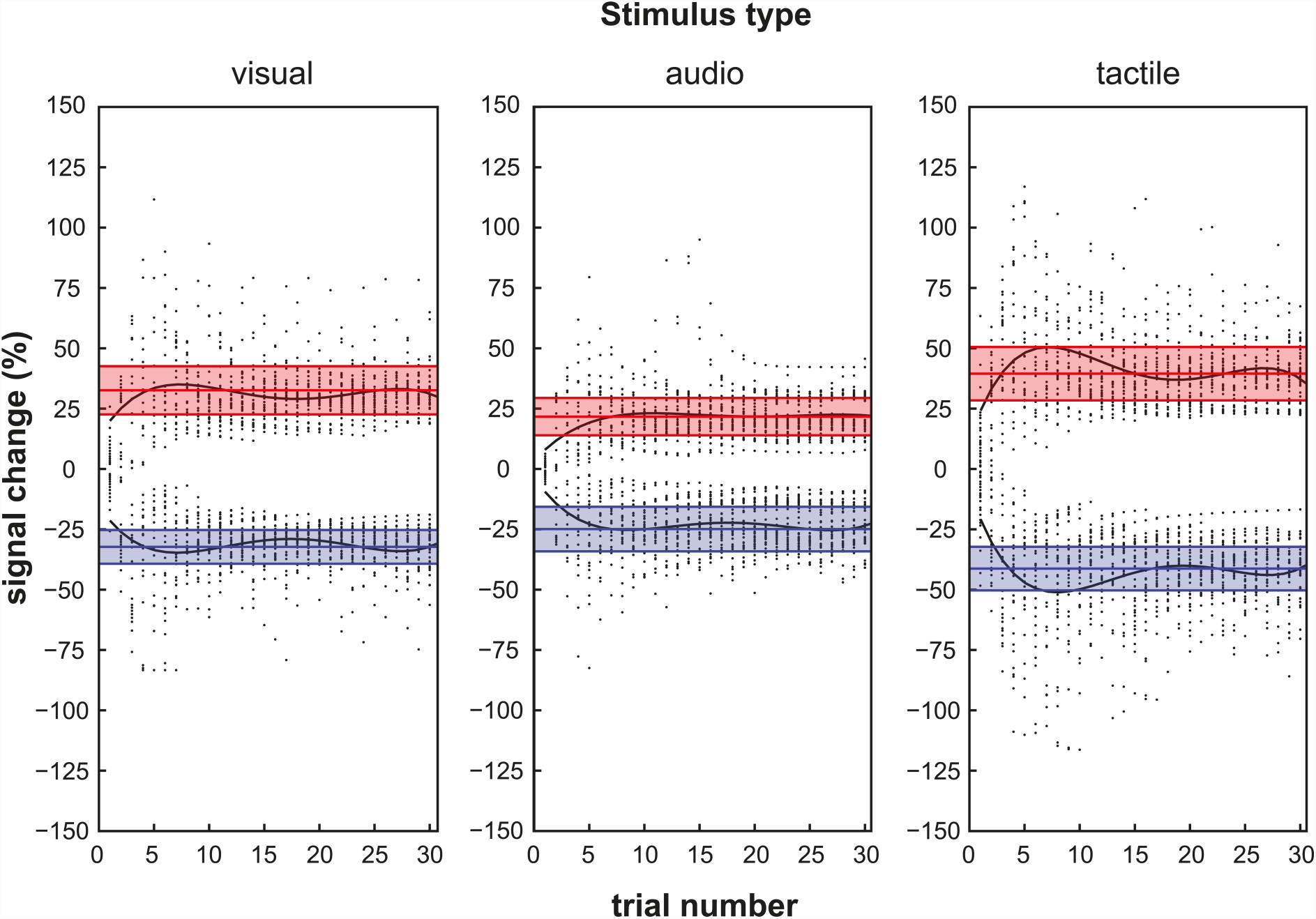
Summary of the thresholding procedure for the detection of visual, auditory and tactile stimulus intensity changes. Scatter plots depict stimulation intensity change (y-axis) per trial (x-axis) from all participants. Black lines depict average progression of the detection threshold estimation (line of best fit for the scatter plot). Colored lines are drawn at the average final threshold (increases in red, decreases in blue; shaded areas represent standard error of the mean).

### 2.5 Analysis

All data processing and analysis was performed with MATLAB (Release 2012b, The MathWorks, Inc., Natick. Massachusetts, United States). Response times (RTs) of correct responses were subject to an outlier analysis that was performed separately for each stimulus configuration, attentional focus and participant. RTs below or above an absolute z-transformed RT of 3 were excluded from further analysis, which was on average 0.87 % of all trials. The statistical analysis was focused on congruence between attended stimuli (*congruence*), the impact of perceptual competition (*distractor*) within the different attentional conditions (*attention*), and respective interactions. The direction of changes per se was ignored. Mean RTs and accuracies (ACCs) were calculated for each level of factors *attention* (VT, AV, AT), *congruence* (attended congruent versus attended incongruent), and *distractor* (congruence relation of unattended stimulus to attended stimuli).

In a 3 (*attention*) x 2 (*congruence*) repeated-measures ANOVA, interactions between *attention* and *congruence* were analysed. Simple effects of *attention* and *congruence* were computed using paired-sample t-tests with an alpha correction according to Holm-Bonferroni (Holm, 1979). The interaction was followed up by computing paired-sample t-tests between congruence differences (congruent – incongruent; noted, for instance, as VT_dif_) for all attentional foci. In order to analyse the impact of *distractor*, two separate 3 (*attention*) x 2 (*distractor*) repeated-measures ANOVA for the levels of the factor *congruence* were computed. This was done because the factor *distractor* is ambiguous with respect to the levels of *congruence*. For attended congruent conditions, distractor differentiates between fully congruent stimuli and a deviant unattended stimulus. For attended incongruent conditions, distractor contrasts the two possible congruence relations of the unattended stimulus with one of the attended stimuli. Simple effects of *distractor* were analysed further based on normalized difference scores. The net difference between *distractor* levels within a given attentional focus was divided by the mean of both values and scaled by 100. These normalized differences were subjected to one-sample t-tests against 0, interactions were evaluated with paired-sample t-tests between these differences (again, noted as for instance VT_dif_; alpha was adjusted according to Holm-Bonferroni). If sphericity was not given, reported values are corrected according to Greenhouse-Geisser. Effect sizes are given as partial eta squared (η^2^). A priori estimation of required sample size was carried out for the 2 x 3 ANOVA reported above using G*Power (Faul, Erdfelder, Lang, & Buchner, 2007). Alpha error probability was set to 5 % and power to 95 %. For detecting a medium sized interaction effect (f = 0.3, cf. Cohen, 1988), calculations suggested a sample size of N = 31. No stopping rule was used.

Complementary to ANOVA, vincentized cumulative distribution functions (vCDFs) were computed and analysed using the same contrasts as for the ANOVA. Per subject, RTs per subcondition were sorted and 21 quantiles were extracted. The resulting vCDFs were fitted to Gamma functions by computing maximum likelihood estimates of shape and scale parameters. Based on the estimated parameters, continuous CDFs were computed. T-tests of contrasts were computed for bins of 5 ms between 500 ms and 3000 ms of the continuous CDFs and p-values were corrected for multiple comparisons by false discovery rate correction (α = .05).

## 3 Results

The results of the thresholding procedure are outlined in Figure 3. Smallest detection thresholds were found for auditory changes (mean percentage deviation from baseline intensity with standard deviations; increase: 21.64 ± 7.74, decrease: 24.93 ± 9.22), medium-sized thresholds for visual changes (increase: 32.60 ± 9.98, decrease: 32.28 ± 6.99) and largest thresholds for tactile changes (increase: 39.59 ± 11.08, decrease: 41.28 ± 9.03). Average response times and response accuracies are depicted in Table 1.

### 3.1 Attention and Congruence

#### Response times

The *attention* x *congruence* ANOVA (aggregating across levels of *distractor*) revealed significant main effects of *attention* and *congruence*, as well as a significant interaction (Fig. 4, left panel; *attention*: *F*_2,66_ = 27.76, *p* < .001, η^2^ = .45; *congruence*: *F*_1,33_ = 50.06, *p* < .001, η^2^ = .60; interaction: *F*_2,66_ = 25.30, *p* < .001; η^2^ = .43). Participants gave fastest responses in condition VT and were slowest in AT (VT–AV: t_33_ = – 3.31, *p* = .002; AV–AT: t_33_ = –4.43, *p* < .001; VT–AT: t_33_ = –6.24 *p* < .001). Congruent responses were faster than incongruent responses in all attention conditions (VT: t_33_ = –3.55, *p* = .001; AV: t_33_ = –3.97, *p* < .001; AT: t_33_ = –10.48, *p* < .001). The difference between attended congruent and attended incongruent conditions varied with respect to the levels of *attention*. While this difference was of comparable in size between VT and AV (AV_dif_ – VT_dif_: t_33_ = –1.54, *p* = .134), AT showed a considerably larger difference compared to both other attentional foci (AV_dif_ – AT_dif_: t_33_ = –4.73, *p* < .001; VT_dif_ – AT_dif_: t_33_ = –6.33, *p* < .001).

**Fig. 4.**
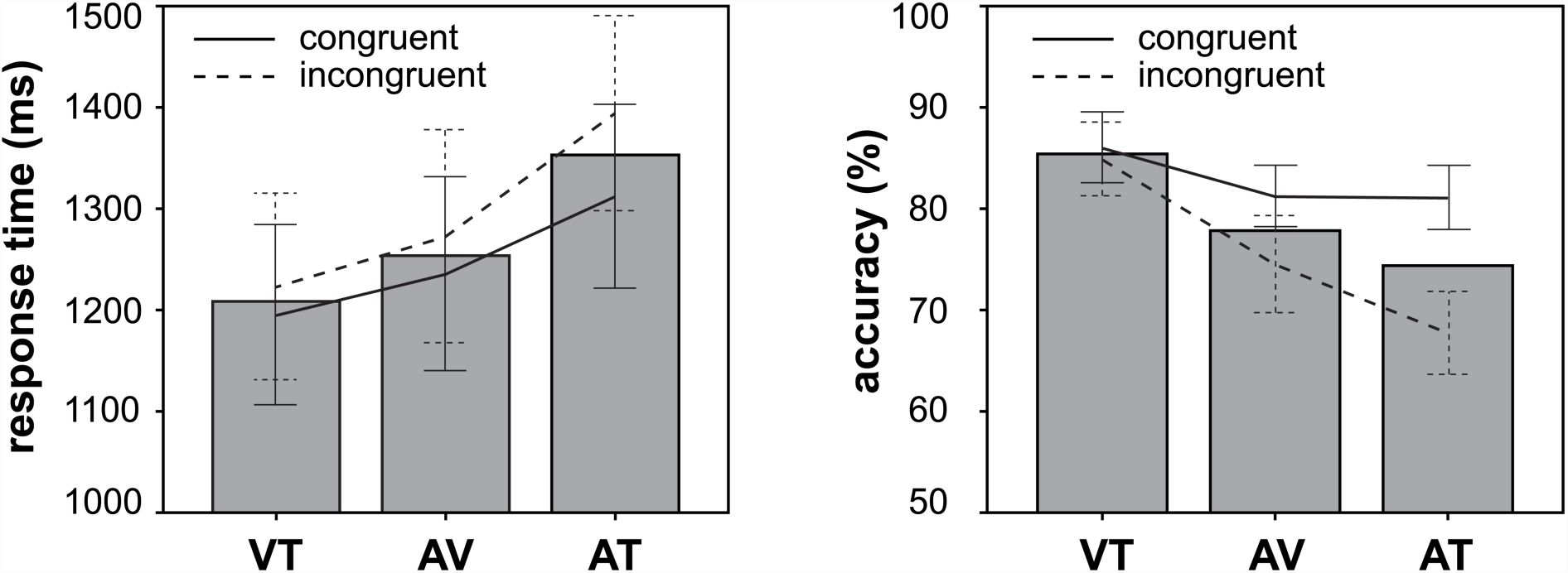
Response times (left panel) and accuracy (right panel) by factors *attention* and *congruence*. Bars depict mean values per levels of *attention* (VT = visual-tactile, AV = audio-visual, AT = audio-tactile). Lines depict differences within attentional conditions with respect to congruence between attended modalities (error bars represent 95 % confidence intervals).

**Table 1.** Mean response times (RT) and accuracies (ACC)

|  | T=A=V | V=T≠A | A=V≠T | A=T≠V |
| --- | --- | --- | --- | --- |
| VT |  |  |  |  |
| ACC | 88.97 | 82.98 | 85.48 | 84.19 |
|  | [85.89, 92.05] | [78.66, 87.30] | [81.59, 89.36] | [80.43, 87.95] |
| RT | 1181.03 | 1208.03 | 1222.01 | 1222.97 |
|  | [1091.04, 1271.02] | [1118.86, 1297.20] | [1126.07, 1317.96] | [1133.80, 1312.14] |
| AV |  |  |  |  |
| ACC | 86.25 | 69.93 | 76.14 | 79.04 |
|  | [83.55, 88.95] | [64.33, 75.52] | [72.09, 80.19] | [74.68, 83.41] |
| RT | 1224.27 | 1281.76 | 1246.06 | 1262.87 |
|  | [1126.19, 1322.34] | [1173.24, 1390.28] | [1151.14, 1340.98] | [1159.87, 1365.88] |
| AT |  |  |  |  |
| ACC | 82.72 | 65.88 | 69.56 | 79.38 |
|  | [97.58, 85.86] | [61.71, 70.05] | [65.20, 73.92] | [75.68, 83.07] |
| RT | 1299.64 | 1393.49 | 1395.02 | 1324.24 |
|  | [1210.47, 1388.81] | [1298.35, 1488.63] | [1296.39, 1493.66] | [1230.54, 1417.94] |

Note: VT = visual-tactile, AV = audio-visual, AT = audio-tactile; ‘=’ denotes ‘is congruent to’, ‘≠’ denotes ‘is incongruent to’. Accordingly, ‘V=T≠A’ specifies trials in which visual and tactile stimuli changed congruently but the tactile stimulus was incongruent to both other stimuli; 95 % confidence intervals are given in brackets below the mean values.

#### Accuracy

With respect to the accuracy of responses, the same overall picture of results emerged as for RTs (Fig. 4, right panel; *attention*: *F*_2,66_ = 27.25, *p* < .001, η^2^ = .45; *congruence*: *F*_1,33_ = 44.06, *p* < .001, η^2^ = .57; interaction: *F*_2,66_ = 21.84, *p* < .001; η^2^ = .40). Participants were most accurate in VT and made most errors in AT (VT – AV: t_33_ = –5.14, *p* < .001; AV – AT: t_33_ = –2.45, *p* = .020; VT – AT: t_33_ = –6.55 *p* < .001). On average, participants made significantly less errors in attended congruent conditions. In contrast to RTs, participants’ accuracy in VT was not better for attended congruent compared to attended incongruent conditions (VT: t_33_ = 1.19, *p* = .245). Still, significant differences between attended congruent and attended incongruent conditions were found for AT and AV (AV: t_33_ = 4.27, *p* < .001; AT: t_33_ = 7.23, *p* < .001). This difference was larger in AT (AV_dif_ – AT_dif_: t_33_ = 3.30, *p* = .002).

#### CDFs

Analysis of CDFs showed a similar pattern of results. Significant differences between attentional foci were found in all three comparisons (Fig. 5a). Cumulative probability of response times (cpRT) in VT was significantly larger in the interval [1500,1800] ms compared to AV. In an interval of [1000,2000] ms, AV showed significantly larger cpRT compared to AT. The largest interval of differences in cpRT was found between VT and AT with [900,2100] ms. Comparisons of CDFs between attended congruent and attended incongruent conditions also matched the result pattern from ANOVA (Fig. 5b). For VT, cpRT in an early interval of [500,1400] ms was significantly larger for attended congruent compared to attended incongruent conditions. Comparably, cpRT in AV differed with respect to congruence in an interval of [800,1500] ms. For AT, cpRT was larger for attended congruent conditions for almost the entire response interval of [500,2600] ms.

**Fig. 5.**
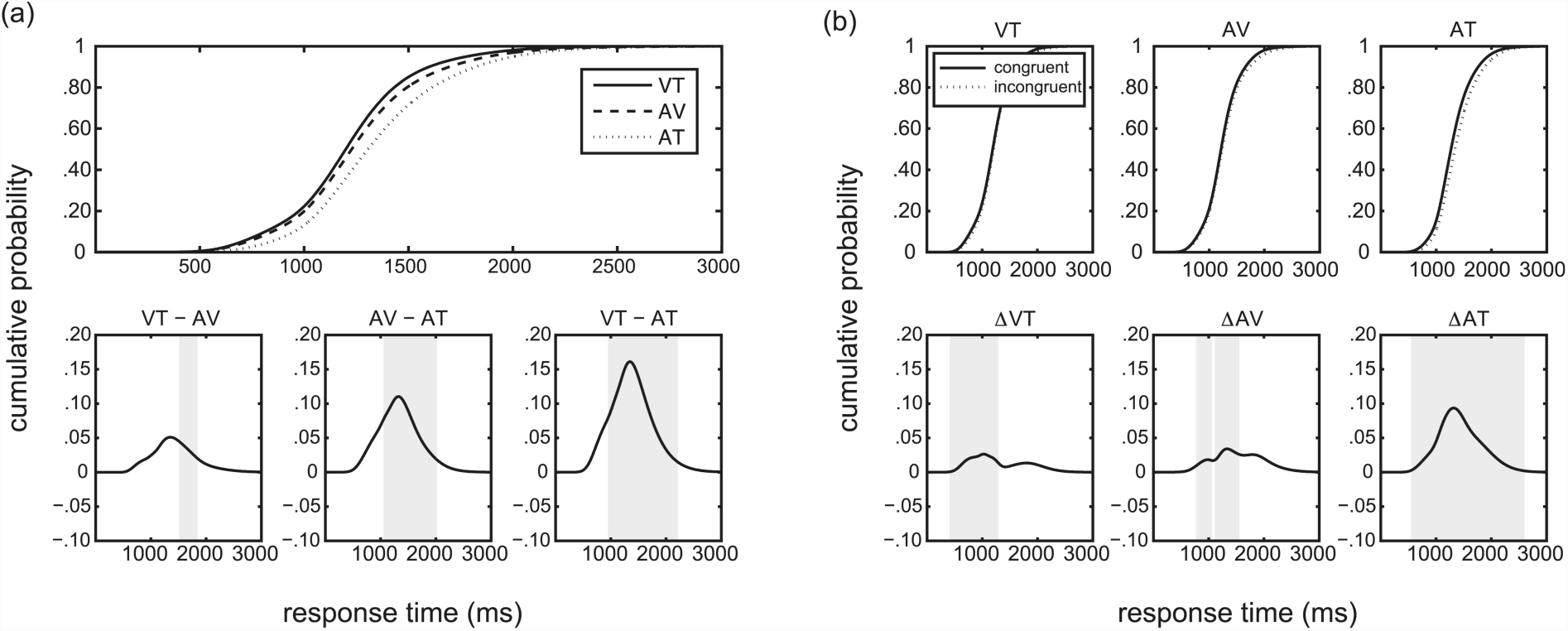
Cumulative density functions (CDFs) of RTs (upper row) and differences between CDFs (lower row). Grey shaded areas in the bottom panels represent regions where contrasted CDFs differ significantly (p < .05, FDR corrected at α = .05). (a) CDFs represent grand means per attentional focus, i.e. visual–tactile (VT), audio–visual (AV), and audio–tactile (AT). The bottom panel shows the differences between the CDFs. (b) CDFs represent grand means of attended congruent respectively attended incongruent conditions per attentional focus. Differences in the lower panel are congruent – incongruent.

### 3.2 Attention and Distraction

#### 3.2.1 Attended congruent

In the following two paragraphs, results from the *attention x distractor* ANOVA on response times and accuracies from attended congruent conditions will be reported (Fig. 6, left column). The factor *distractor* contrasts between a converging (fully congruent, Fig. 1, I.a) and deviant irrelevant modality (congruent with distraction, Fig. 1, I.b). In addition, statistical analyses on CDFs employing the same contrasts will be reported (Fig. 7a).

**Fig. 6.**
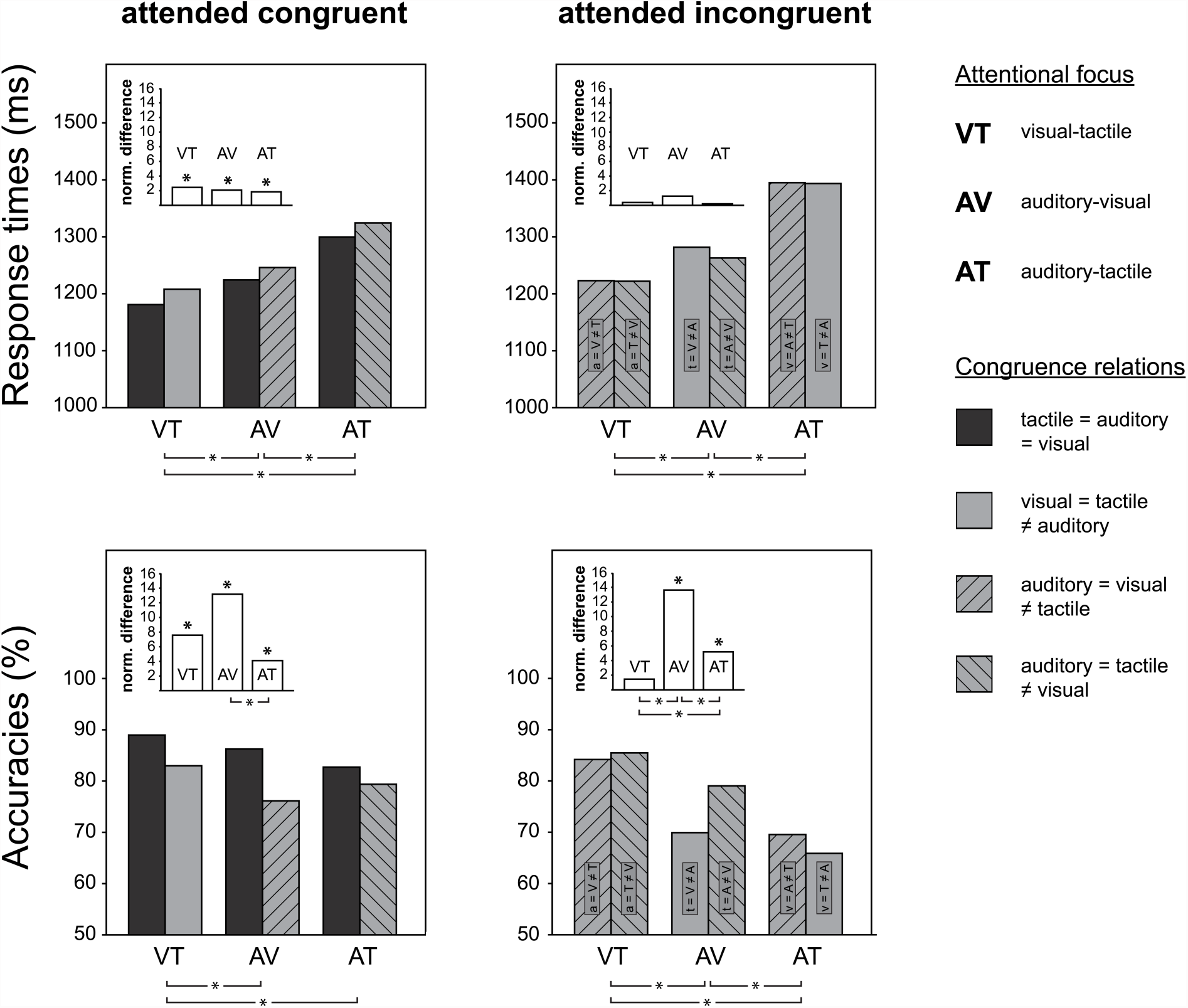
Response times and accuracies as function of factors *attention* and *distraction.* Bars depict averages of response times (upper row) and accuracies (lower row). For attended congruent, the two bars per attentional condition depict a congruent respectively incongruent distractor (left column). For attended incongruent, the two bars per attentional condition depict different congruence relations of the distractor to one or the other attended stimulus (right column; capital letters in bar labels indicate attended modalities, small letters the unattended modality). Color-coding indicates the congruence pattern of the trimodal stimulus (legend on right margin). Significant mean differences are indicated by an asterisk below each plot. The subplots depict absolute values of normalized differences (see Method section for details) between *distractor* levels (neighbouring bars per attentional focus). Again, asterisks indicate significant differences tested against 0 (above bars) and significant differences of these values across attentional conditions (below chart). Alpha levels were adjusted according to Holm-Bonferroni. Note for the legend: ‘=’ denotes ‘is congruent to’, ‘≠’ denotes ‘is incongruent to’.

**Fig. 7.**
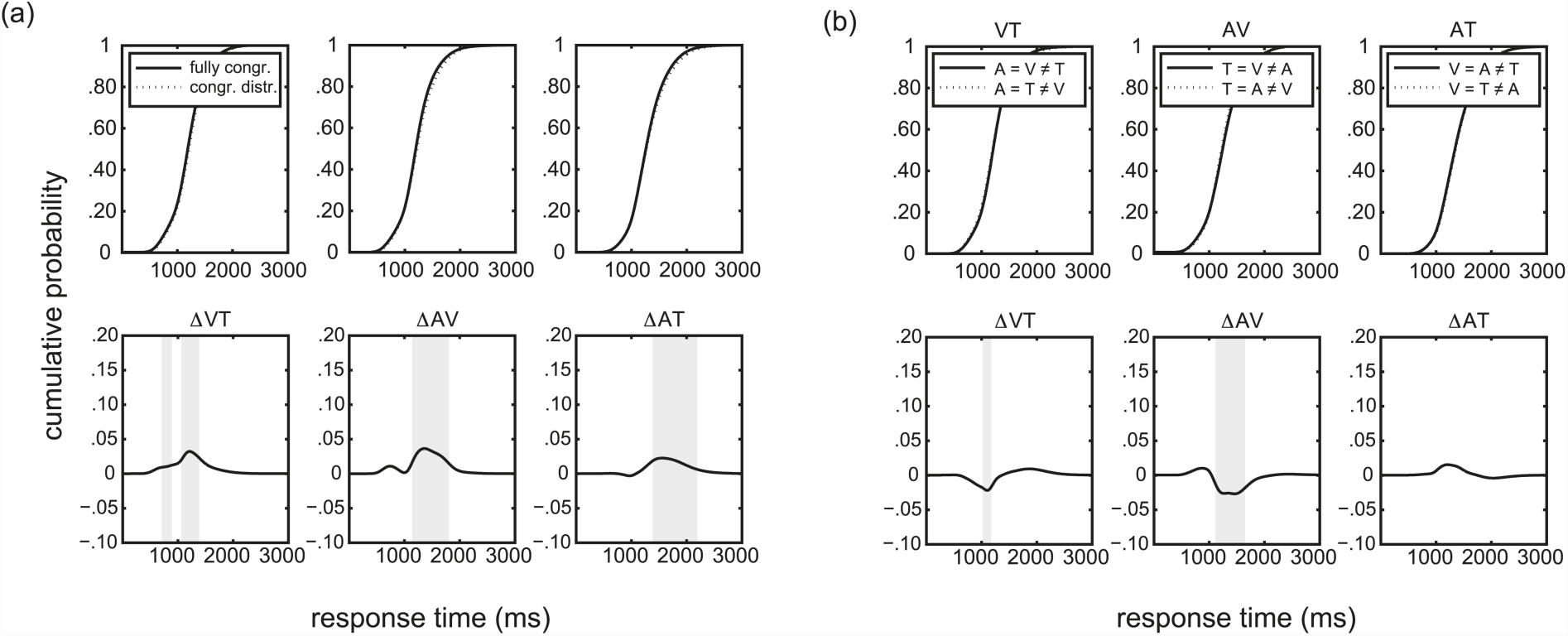
Cumulative density functions (CDFs) of RTs (top panels) and differences between CDFs (bottom panels). Grey shaded areas in the bottom panels represent regions where contrasted CDFs differ significantly (p < .05, FDR corrected at α = .05). (a) CDFs represent grand means of fully congruent (fully congr.) respectively congruent distracted (congr. distr.) conditions per attentional focus. Differences in the bottom panel are fully congruent – congruent distracted. (b) CDFs represent grand means of the two possible congruence relations of the respective distractor to one or the other attended modalities. Differences in the bottom panel are solid line – dotted line.

##### Response times

Participants’ response times were significantly affected by *attention* and *distractor,* but not their interaction (Fig. 6. upper left panel; *attention*: *F*_2,66_ = 18.53, *p* < .001, η^2^ = .36; *distractor*: *F*_1,33_ = 29.42, *p* < .001, η^2^ = .47; interaction: *F*_2,66_ = .10, *p* > .250). Again, participants gave fastest responses in condition VT and were slowest in AT (VT – AV: t_33_ = – 2.99, *p* = .005; AV – AT: t_33_ = –3.67, *p* = .001; VT – AT: t_33_ = –5.10 *p* < .001). Normalized differences across distractor levels indicated that the presence of a diverging distractor led to a significant slowing of participants’ responses in all attentional foci (VT: t_33_ = –3.97, *p* < .001; AV: t_33_ = –3.29, *p* = .002; AT: t_33_ = –3.05, *p* = .004).

##### Accuracy

Accuracy of responses was significantly modulated by *attention*, *distractor*, and the interaction effect (*attention*: *F*_2,66_ = 6.38, *p* = .004, η^2^ = .16; *distractor*: *F*_1,33_ = 63.99, *p* < .001, η^2^ = .66; interaction: *F*_2,66_ = 5.41, *p* = .007, η^2^ = .14). Participants made significantly more errors if the distractor diverged from the congruent attended modalities in all attentional foci (VT: t_33_ = 4.07, *p* < .001; AV: t_33_ = 6.08, *p* < .001; AT: t_33_ = 2.63, *p* = .013). Participants were significantly more accurate in VT than in both AV and AT (VT – AV: t_33_ = 3.51, *p* = .001; VT – AT: t_33_ = 2.72, *p* = .010), but performance in AV and AT was comparable (AV – AT: t_33_ = 0.10, *p* > .250). While participants were equally strong distracted in VT and AV as well as VT and AT (VT_dif_ – AV_dif_: *p* = .059, VT_dif_ – AT_dif_: *p* = .219), tactile divergence in AV distracted significantly stronger than visual divergence in AT (t_33_ = 3.09, *p* = .004).

##### CDFs

Again, results from the analysis of CDFs largely mirrored the results from ANOVA. Significant differences between fully congruent and congruent distracted conditions were found for all attentional foci (Fig. 7a). In VT, an incongruent distractor was associated with significantly lower cpRT in an interval of [750,1400] ms. In an interval of [1100,1800] ms, cpRT in AV were significantly lower if the distractor was incongruent. In AT, this difference was significant in an interval of [1500,2000] ms. Despite the differences between the attentional foci with respect to the segment of the CDFs where significant differences were found, the length of intervals was comparable. This parallels the comparable magnitudes of mean differences in RTs found in the ANOVA.

#### 3.2.2 Attended incongruent

Figure 6 (right column) illustrates the results concerning the attended incongruent condition. In the respective *attention* x *distrator* ANOVA, the factor *distractor* contrasts between congruence relations of the respective distractor with one or the other attended modality (e.g. the difference between performance on Fig. 1., II.a+b). The corresponding CDFs are reported in Figure 7.b.

##### Response times

The timing of responses differed significantly with respect to *attention* (upper right panel of Fig. 6; *F*_2,66_ = 33.41, *p* < .001, η^2^ = .36) but neither *distractor* nor the interaction significantly affected RTs (*distractor*: *F*_2,66_ = 2.24, *p* = .144; interaction: *F*_1,33_ = 2.95, *p* = .069). Again, participants were fastest in VT and slowest in AT (VT – AV: t_33_ = – 3.46, *p* = .002; AV – AT: t_33_ = –4.92, *p* < .001; VT – AT: t_33_ = –7.12, *p* < .001).

##### Accuracy

*Attention* affected participants’ accuracy most strongly (lower right panel of Fig. 6; *F*_2,66_ = 38.05, *p* < .001, η^2^ = .54). Participants showed highest accuracy in VT and made most errors in AT (VT – AV: t_33_ = 5.48, *p* < .001; AV – AT: t_33_ = 3.50, *p* = .001; VT – AT: t_33_ = 8.15 *p* < .001). Further, *distractor* and the interaction affected the accuracy of responding (*distractor*: *F*_1,33_ = 8.75, *p* = .006, η^2^ = .21; interaction: *F*_2,66_ = 23.71, *p* < .001, η^2^ = .42). Participants were significantly less accurate in AV when the tactile distractor was congruent to the visual compared to when congruent to the auditory target (t_33_ = 5.83, *p* < .001). Likewise, accuracy was lower in AT when the visual distractor was congruent to the tactile target compared to when congruent to the auditory target (t_33_ = 2.67, *p* = .012). In VT, participants responded equally accurate in both conditions (*p* > .250). Thus, the differences in AV and AT were significantly larger than in VT (AV_dif_ – VT_dif_: t_33_ = 3.99, *p* < .001; AT_dif_ – VT_dif_: t_33_ = 2.41 *p* = .022). In addition, the difference in AV was significantly larger than in AT (AV_dif_ – AT_dif_: t_33_ = 7.40, *p* < .001).

##### CDFs

While no effects of the distractor were found in the analysis of mean RTs of the attended incongruent conditions (see above), significant differences were found in the analysis of the CDFs. In VT, cpRT was significantly lower if the auditory distractor was congruent to the attended visual stimulus compared to the condition where the distractor was congruent to the attended tactile stimulus in a narrow interval of [1000,1100] ms. In AV, significant differences with respect to the congruence relation of the distractor were found in an interval of [1100,1600] ms. In that segment of CDFs, cpRT was significantly lower if the tactile distractor was congruent to the attended visual stimulus compared to the case where the distractor was congruent to the attended auditory stimulus. For AT, no differences with respect to the congruence relation of the distractor were found.

## 4 Discussion

We investigated differences in crossmodal matching between bimodal combinations of visual, auditory, and somatosensory stimuli. Crossmodal congruence was used to modulate stimulus-driven mechanisms of multisensory integration. In line with our expectations, we found better performance for vision-including attentional foci. Furthermore, we observed that congruence between attended stimuli was associated with better performance compared to incongruent conditions. The difference between attended congruent and attended incongruent conditions was inversely related to the overall level of performance in the attentional foci. That is, the largest difference between congruent and incongruent conditions was found for audio-tactile matching which overall showed longest RTs and lowest accuracy. Conversely, performance for visual-tactile matching was best overall and showed the smallest difference between congruent and incongruent conditions.

### 4.1 Performance differences between bimodal foci

Clear mean differences between attentional foci were found. Not only vision-including foci (AV and VT) showed better performance than the audio–tactile focus, but also VT compared to AV. These differences cannot be explained by salience differences between the modalities because individual thresholds for each stimulus were estimated elaborately and balanced across modalities. Our hypotheses were based on the assumption that performance in crossmodal matching would most critically depend on the ability to inhibit the distracting modality. In addition to that, perceptual gains of the modalities to be matched might influence crossmodal integration. Wang et al (2012) showed that perceptual competition differentially modulated perceptual gains of visual, auditory, and somatosensory stimuli. Accordingly, performance in visual–tactile matching might have been best because the tactile component of the multisensory stimulus was enhanced and the visual component remained unchanged with respect to perceptual gain. Conversely, audio–tactile integration might have been more difficult because the tactile component was enhanced while the auditory component was suppressed. This pattern would suggest that best performance in crossmodal matching is achieved if the modalities to be integrated have comparable levels of perceptual gain. A recent modeling approach of audio-visual redundant signal detection led to a similar conclusion and postulated that crossmodal integration can be expected to be most beneficial for signals that would individually lead to comparable performance levels (Otto et al., 2013).

### 4.2 Crossmodal congruence and perceptual competition

Crossmodal congruence between the task-irrelevant modality and attended congruent modalities improved performance in all attentional foci with comparable magnitude. This boost in performance by irrelevant but congruent sensory information suggests that crossmodal congruence enhances perceptual processing irrespective of attention. Alternatively, this performance difference might merely be related to the absence of conflicting information in the fully congruent trials. Evidence from attended incongruent conditions, however, supports the notion that crossmodal congruence affects perceptual processing directly. Under the assumption that perceptual competition by the distractor differs only with respect to it being conflicting or not, performance in attended incongruent conditions should not differ with respect to the congruence relation of a given distractor to one or the other attended modalities. This, however, was the case and produced the inverse relationship between overall performance and differences between congruent and incongruent conditions per attentional focus.

For the audio–visual focus, attended incongruent performance was significantly lower if the tactile distractor was congruent to the attended visual modality compared to when it was congruent to the attended auditory modality ([*t* = *V*] < [*t* = *A*])^1^. Similarly, attended incongruent performance for the audio–tactile focus was lower when the visual distractor was congruent to the attended tactile modality compared to when it was congruent to the attended auditory modality ([*v* = *T*] < [*v* = *A*]). For the visual–tactile focus, however, there was no difference in performance with respect to the congruence relation of the distractor to the attended modalities ([a = V] = [a = T]). The resulting pattern of performance (i.e., [*V* = *T*] < [*A* = *V*]) ≤ [*A* = *T*]) mirrors the pattern of mean performance for the attentional foci noted earlier. We conclude that crossmodal congruence enhancement is most effective for crossmodal pairs that have comparable levels of perceptual gain. Therefore, perceptual processing of visual–tactile stimulus combinations is enhanced most strongly, but less strongly for both audio–visual and audio–tactile pairs.

### 4.3 Multisensory integration and processing stages

Crossmodal interactions can take place at multiple stages of information processing and at various cortical sites (Ghazanfar & Schroeder, 2006). While phenomena of integration can be observed already in primary sensory areas, more complex interactions occur at superior temporal or posterior parietal as well as prefrontal cortical areas (Werner & Noppeney, 2010). With respect to the current results, it is unclear at what stages of processing crossmodal interactions occurred. We speculate that early inter-sensory crosstalk (Talsma et al., 2010) might have led to adaptations in perceptual gain as described above. In subsequent processing, mechanisms of crossmodal matching and congruence enhancement were constrained by the gain adjusted multisensory input. In order to test these assumptions, electrophysiological recordings might be best suited to investigate the dynamics of neural processing in this paradigm. Early sensory processing should reflect the aforementioned gain adjustments and thus should not differ between attentional conditions. Congruence relations should systematically affect cortical processing at later stages of processing, for instance in temporal, parietal and/or frontal regions (Göschl, Friese, Daume, König, & Engel, 2015).

## 5 Conclusions

Employing a novel trimodal paradigm, we found that crossmodal congruence improved performance if both, the attended modalities and the task-irrelevant distractor were congruent. On the other hand, if the attended modalities were incongruent, the distractor impaired performance because of its congruence relation to one of the attended modalities. Whether this effect is based on an automatic relocation of attention to the congruent stimulus pair or due to a conflict to the required response, could not be decided in this study. Between attentional conditions, magnitudes of crossmodal enhancement or respectively impairment differed. Our results suggest that crossmodal congruence has the strongest effects on visual– tactile pairs, intermediate effects for audio–visual pairs and weakest effects for audio–tactile pairs. The origin of such differences might be evolutionary. That is, statistical properties of the environment may have resulted in use-dependent differentiation of crossmodal mechanisms. Vision and touch, for instance, often occur congruently, in many cases even predictably, e.g., if self-initiated movements of the hands trigger visual–tactile percepts. In contrast, audio–visual as well as audio–tactile stimulus combinations predominantly have multiple external sources and are thus less reliable. This graded pattern of crossmodal interactions might relate to different patterns of functional connectivity in multisensory neural networks, and possibly even to differences in structural connectivity between sensory cortices. In addition to electrophysiological studies on dynamic functional coupling as a basis of crossmodal integration, structural imaging could contribute to revealing variations across different multisensory networks.

## Author contributions

A. K. Engel developed the study concept. All authors contributed to the study design. J. Misselhorn performed testing and data collection. J. Misselhorn conducted data analysis and interpretation with advice from U. Friese under supervision of A. K. Engel. J. Misselhorn drafted the manuscript, and all other authors provided critical revisions. All authors approved the final version of the manuscript for submission.

## Acknowledgments

This research was supported by grants from the German Research Foundation (SFB 936/A3) and the European Union (ERC-2010-AdG-269716).

## Footnotes

1 Capital letters indicate attended modalities and small letters indicate unattended modalities.

